# Circadian pacemaker neurons display co-phasic rhythms in basal calcium level and in fast calcium fluctuations

**DOI:** 10.1101/2021.05.03.442479

**Authors:** Xitong Liang, Timothy E. Holy, Paul H. Taghert

## Abstract

Circadian pacemaker neurons in the *Drosophila* brain display daily rhythms in the levels of intracellular calcium. These calcium rhythms are driven by molecular clocks and are required for normal circadian behavior. To study their biological basis, we employed genetic manipulations in conjunction with in vivo light-sheet microscopy to measure calcium dynamics in individual pacemaker neurons over complete 24-hour periods. We found co-phasic daily rhythms in basal calcium levels and in high frequency calcium fluctuations. Further we found that the rhythms of basal calcium levels require the activity of the *IP3R*, a channel that mediates calcium fluxes from internal endoplasmic reticulum (ER) calcium stores. Independently, the rhythms of fast calcium fluctuations required the T-type voltage-gated calcium channel, a conductance that mediates extracellular calcium influx. These results suggest that *Drosophila* molecular clocks regulate *IP3R* and T-type channels to generate coupled rhythms in basal calcium and in fast calcium fluctuations, respectively. We propose that both internal and external calcium fluxes are essential for circadian pacemaker neurons to provide rhythmic outputs, and thereby regulate the activities of downstream brain centers.

## Introduction

Circadian rhythms in multiple aspects of cellular physiology help organisms across taxa, from unicellular cyanobacteria to multicellular animals, adapt to environmental day-night changes (Dunlap 1999; Herzog 2007). In mammals, neurons in the hypothalamic suprachiasmatic nucleus (SCN) show circadian rhythms in gene expression, intracellular calcium, neural activity, and other cellular properties (Welsh et al 2010). Circadian rhythms in SCN neuronal outputs coordinate circadian rhythms in other cells throughout the body and generate behavioral rhythms (Mohawk et al 2012). The rhythms of SCN neuronal outputs can be generated cell-intrinsically by the negative transcription/translation feedback loop of core clock genes, as a molecular clock, which then generates 24-hour oscillations in a series of genes (Welsh et al 1995; Dunlap 1999; Panda et al 2002; Colwell 2011). These gene oscillations then regulate different aspects of membrane physiology, such as the expression levels of channels for potassium (K^+^), sodium (Na^+^), and calcium (Ca^2+^) (Pennartz et al 2002; Itri et al 2005; Pitts et al 2006; Meredith et al 2006; Flourakis et al 2015). The mechanisms by which the molecular clockworks coordinate complex membrane physiology to generate neural activity rhythms within individual circadian pacemakers remain to be defined.

Calcium signaling regulates many cellular processes, such as neural excitability, neurotransmitter release, and gene expression (Berridge 1998). Cytoplasmic calcium can be regulated from extracellular calcium influx, as well as from intracellular calcium stored in the endoplasmic reticulum (ER) and mitochondria (Chorna and Hasan 2012). Studies on SCN neurons *in vitro* (Colwell 2000; Ikeda et al 2003) and recently *in vivo* (Jones et al 2018) measured circadian calcium rhythms in SCN neurons. Some studies suggested that calcium rhythms were driven by neuronal firing and voltage-gated calcium channels (Colwell 2000; Enoki et al 2017), while others suggested they were driven by intracellular stores, via the ER channels ryanodine receptor (Ikeda et al 2003). These alternative hypotheses may derive from the technical differences in the various studies, including the details of *in vitro* preparations, but also due to a lack of single-cell resolution in the calcium measurements.

In *Drosophila*, circadian pacemaker neurons also show clock-driven circadian calcium rhythms (Liang et al 2016). The dynamics can be resolved across all five major pacemaker groups, s-LNv, l-LNv, LNd, DN1, and DN3, and each group exhibits distinct and sequential daily peak phases. Within such groups, the rhythms can be measured in single identified cells (Liang et al., 2017). The multi-hour phase diversity exhibited by this network requires a series of delays effected by environmental light and by non-cell-autonomous modulation mediated by different neuropeptides (Liang et al 2017). Precisely how neuropeptide signaling regulates calcium activity in pacemaker neurons over long (many-hour) durations is unknown. To begin to understand these critical mechanisms of pacemaker modulation, we begin by addressing the cellular and molecular basis of pacemaker calcium rhythms with physiological, genetic and behavioral measures.

In this study, we again used *in vivo* calcium imaging at single-cell resolution, here using a high-speed light sheet microscope (Greer and Holy 2019). We simultaneously measured both basal calcium levels and fast calcium fluctuations over entire 24-hr periods. We found circadian rhythmicity in both basal calcium levels and in the frequency of fast calcium fluctuations. We consider the fast fluctuations to represent events closely coupled to neuronal firing, as have previous, related studies (Pologruto et al 2004, Yaksi and Friedrich 2006, Chen et al., 2013, Streit et al. 2016, Greenberg et al 2018). In all pacemaker neurons studied, these two layers of calcium rhythms shared the same daily temporal pattern (i.e., they were co-phasic). To gain insights into the mechanism of these patterns, we exploited the fact that in *Drosophila* many calcium channels are encoded by single genes (Chorna and Hasan 2012), and used genetics to study the roles of individual channels in generating daily pacemaker calcium rhythms. Here, we present results of experiments in which we knocked down RNAs encoding different calcium channels selectively in all or a subset of pacemakers. We evaluated the impact of individual channels in setting both slow daily changes in basal calcium levels and in fast fluctuations. Finally, we measured PERIOD staining levels and behavior to determine which channels provide feedback to the molecular clock and which are required for normal circadian output from the pacemaker network.

## Results

### The rhythms of slow and fast calcium activity changes show similar daily patterns

Previously we reported that five major groups of circadian pacemaker neurons each exhibit daily calcium rhythms with distinct phases (Liang et al 2016). These results stand in apparent contrast to descriptions of synchronous daily electrical activity rhythms among three of these groups, the s-LNv, l-LNv, and DN1 (Cao and Nitabach 2008; Sheeba et al 2008; Flourakis et al 2015). The electrical activity rhythms were recorded *ex vivo* from different brains isolated at four to six different time points of the day. In contrast, we measured calcium rhythms *in vivo* by scanning individual flies every 10 min for 24 hours. Because of the close peak phases of calcium rhythms in s-LNv, l-LNv, and DN1 (between late night to mid-morning -Liang et al 2016), and because of the coarse sampling of electrophysiological studies, it is not certain whether calcium and electrical activity patterns are in fact distinct. To help clarify apparent differences in results derived from the two sets of studies, we began by performing short-term, continuous *in vivo* calcium imaging (1 Hz volumetric rate) on fly brains that were exposed acutely before each imaging experiment at five different times of day. We focused on the LNd because this group has a phase of calcium rhythms most distinct from those of the s-LNv, l-LNv, and DN1. In addition, the daily electrophysiological activity pattern of LNd has not previously been reported. We found that ∼0.1 Hz calcium fluctuations peaked at around the same ZT8 -ZT10 at which this pacemaker group shows peak intensity in its daily calcium rhythm (Figure 1A and Figure S1). The time course of “fast” (by circadian standards) calcium fluctuations in the evening suggests they might be caused by the calcium influx during single action potentials or bursts of them (Figure 1A, Yaksi and Friedrich 2006, Chen et al., 2013, Greenberg et al 2018). This result suggested that one or more LNd pacemakers exhibit a daily rhythm in electrical neural activity that is roughly co-phasic with this pacemaker group’s slow daily calcium rhythm.

**Figure 1.**
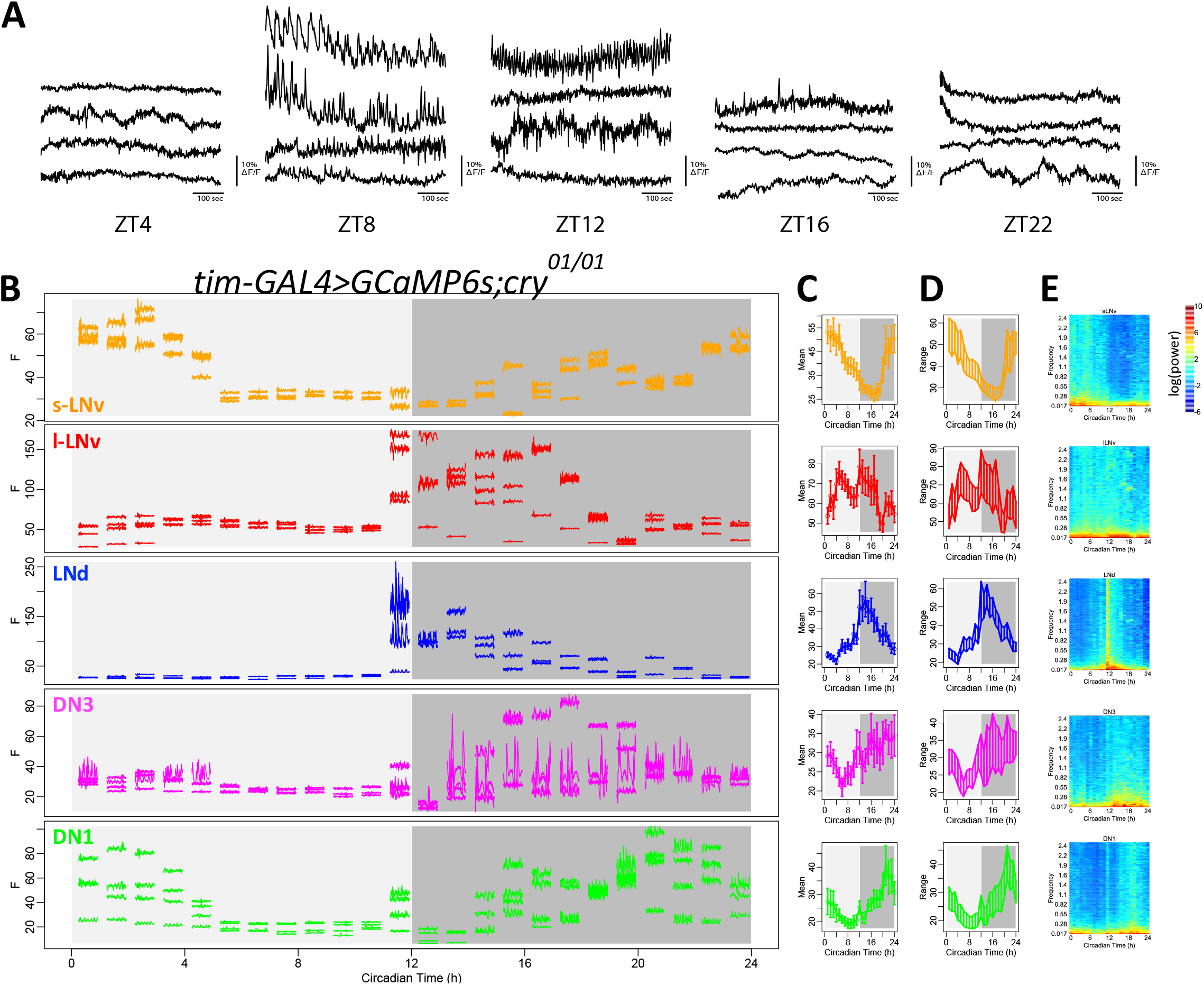
Daily pattern of fast calcium activity in circadian pacemaker neurons. **(A)** Representative calcium activity traces of LNd recorded at 1Hz from five different times of day: ZT4, ZT8, ZT12, ZT16, and ZT22. For each timepoint, the fly’s brains are acutely exposed for short-term imaging. **(B)** Raw calcium activity traces from one representative fly. Each segmented trace is 1-min activity of single neuron recorded at 5Hz. 24 segments are recorded over 24 hours with 1-hour intervals. Activity traces of five circadian pacemaker groups are plotted in separate panels and color-coded. For each group, only cells that can be tracked throughout all 24 recording sessions are showed. In this specific fly, 3 s-LNv, 4 l-LNv, 3 LNd, 4 DN3, and 4 DN1 can be reliably tracked across 24 recording sessions. Circadian pacemaker neurons exhibit daily modulation in basal calcium level and in the frequency of fast calcium spikes. **(C)** Daily pattern of mean calcium intensity over the 1-min recording session at each of the 24 timepoints, averaged across all 6 flies studied. Error bars denote SEM. **(D)** Daily patterns of the range of calcium transients over the 1-min recording session at each of the 24 timepoints, averaged across all 6 flies studied. **(E)** Daily pattern of the power spectrum over the 1-min recording session at each of 24 timepoints, averaged across all 6 flies studied.

Because the slow and fast calcium LNd rhythms are synchronous as measured, it is formally possible that one rhythm is downstream of the other: for example, the slow calcium rhythm could be the consequence of a rhythm in the fast. Alternatively, these two processes could be completely distinct. To better understand the relationships between the two, and better describe their phases across the entire network, we performed a series of short-term (1 min) high-frequency (5 Hz) *in vivo* calcium imaging episodes at 1-hr intervals using the light-sheet microscope OCPI-II (Greer and Holy 2019). In so doing, we tracked both slow basal calcium level and fast calcium fluctuations in the same individual neurons, from all five major circadian pacemaker groups: we collected these data consecutively from single brains for entire 24 hr periods (Figure 1B). To ensure minimal disruption to the circadian clocks due to repeated optical scanning, we used *cry*^01^ flies for these experiments which are null for the internal photoreceptor CRYPTOCHROME. On average, all circadian neuron groups displayed slow calcium rhythms comparable to those we previously reported (Liang et al 2016), except for the l-LNv, which showed a second daily calcium activation right after the time of lights off. Nevertheless, all pacemaker groups displayed daily changes in the minimal calcium level, demonstrating that their basal calcium levels cycle with a daily rhythm (Figure 1CD). We found that within all five pacemaker groups, changes in basal calcium levels and in fast calcium fluctuations shared similar daily patterns: when basal calcium levels were high within a single pacemaker group, that group also exhibited larger-amplitude fast calcium fluctuations. Power spectrum analysis clearly revealed that, for individual neurons within each pacemaker group, calcium activity at all frequency domains increased when the basal calcium level was high (Figure 1E). We asked whether the change in the incidence of high frequency GCaMP6 fluctuations could have a technical basis: specifically, whether it derives from a higher level of photon shot-noise due to the higher baseline intensity. In order to normalize the effect of shot noise, we also calculated the intensity of the calcium signal as the square root of photon number collected from an individual region of interest (ROI). In this analysis, we still found daily rhythms in fast calcium fluctuations (Figure S2). These results support the hypothesis that, in each circadian pacemaker group, fast calcium fluctuations exhibit a daily rhythmic pattern that is co-phasic with a slow daily rhythm in basal calcium levels.

### An RNAi screen to identify potential contributions of different calcium channels

The observations described above support the conclusion that for individual pacemakers, slow and fast calcium activities co-vary across the day. Yet, these observations do not reveal whether the two rhythms are mechanistically linked, or represent independent functions. To identify the sources for different calcium rhythms, and ask about their relatedness, we used RNAi to knockdown different calcium channels. We performed a limited screen for calcium channels, including three subtypes of α1 subunits and one type of α2δ subunits of voltage-gated calcium channels; in addition, we tested two types of store-operated calcium entry (SOCE), *dSTIM* and *dOrai*, and two types of calcium channels on the endoplasmic reticulum (ER): ryanodine receptor (*RyR*) and inositol trisphosphate receptor (*IP3R*); finally, we included the sarco/endoplasmic reticulum calcium-ATPase (*SERCA*). By knocking down these genes selectively in circadian pacemaker neurons, using *tim-GAL4*, or in a subset of eight PDF-positive pacemaker neurons using *pdf-GAL4*, we first tested whether any of these genes are required for normal circadian behavioral rhythms. We found evidence for the involvement of three (Figure 2 and Table 1) as indicated by increases in the percentage of arrhythmic (%AR) flies tested under constant darkness (DD). Reduced expression of a channel on the cytoplasmic membrane, *α1T*, which encodes the α1 subunit for T-type voltage-gated calcium channel, caused the strongest behavioral arrhythmicity when driven by either *pdf-GAL4* or by *tim-GAL4* (65% and 83%) with one of the two RNAi lines tested (KK100082). Likewise, knockdown of expression of the *SERCA* calcium pump, caused strong arrhythmicity in two different RNAi lines. Yet knocking down *SERCA* with the stronger RNAi line in all circadian pacemakers by *tim-GAL4* also shortened the flies’ lifespans: 69% flies died during behavioral experiments. Knockdown of another calcium channel on the ER membrane, *IP3R*, also affected the circadian rhythm in behavior when driven by *tim-GAL4*. These behavioral deficits suggested that *α1T, SERCA*, and *IP3R* might be involved in the regulation of calcium rhythms in circadian pacemaker neurons.

**Figure 2.**
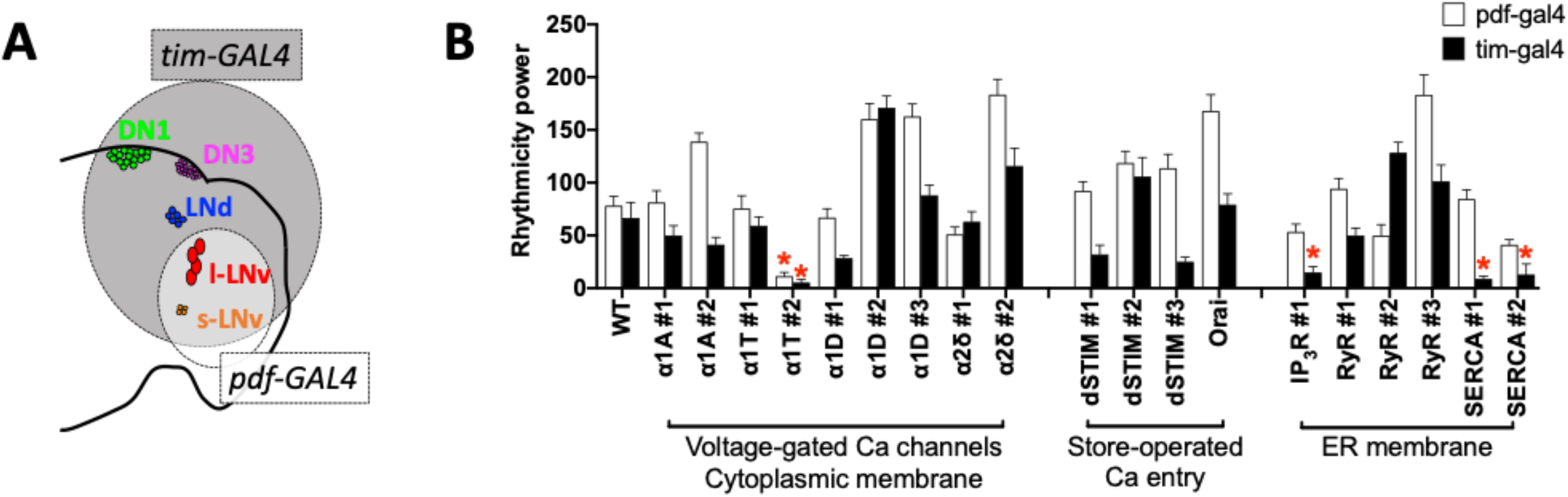
RNAi-screening for calcium channels required for circadian rhythms. **(A)** Schematic of the five pacemaker groups in the brain superimposed with a Venn diagram of expression driven by tim-GAL4 (all groups) and pdf-GAL4 (s- and l-LNV groups). **(B)** Summary of behavioral screening for calcium-channel knockdowns that reduced average rhythm strength in locomotor activity under DD (*p<0.05, two-way ANOVA followed by Dunnett’s multiple comparisons test; also see Table 1).

**Table 1.**
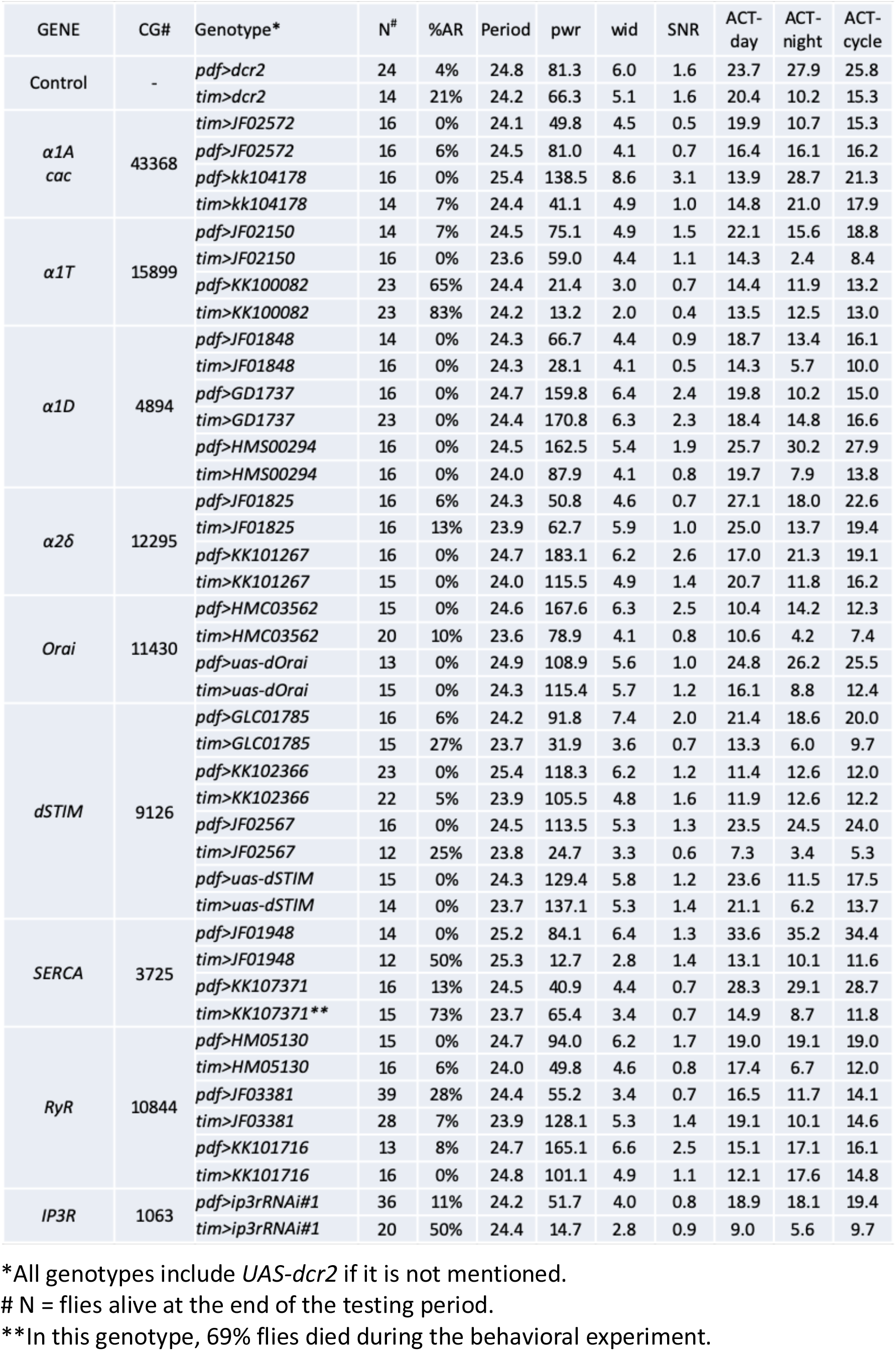
The rhythm strength and period of locomotor activity under constant darkness of flies expressing calcium channel RNAi transgenes in PDF neurons or all clock neurons.

### Slow calcium rhythms require IP3R

We then asked whether the *α1T, SERCA*, and *IP3R* regulating circadian behavior also influence calcium rhythms. We measured GCaMP6 fluorescence during *in vivo* 24-hr recordings in *Drosophila* knock-downs in all, or in just the subset of PDF-positive, circadian neurons (Figure 3A-H). Although knocking down *α1T* caused the strongest behavioral deficits, the slow calcium rhythms of all pacemaker neuron groups in these flies were similar to those in the control genotypes (Figure 3A-D). The amplitude of calcium rhythms in flies with *α1T* knocked down in all pacemaker neurons showed a non-significant trend of decrease to 59.3% on average, while their activity phases were still normal (Figure S3). In contrast, when *SERCA* was knocked down in PDF neurons (Figure 3E), or in all circadian neurons (Figure 3F, using the stronger RNAi line KK107371), the slow calcium activities of these neurons were largely arrhythmic. The amplitudes of calcium fluctuations were decreased to 37.8% on average (Figure S3), and the coherence was lost within groups (Rayleigh test, P > 0.2). Likewise, the calcium rhythms were still normal when *IP3R* was knocked down in PDF neurons (Figure 3G) but became largely arrhythmic when *IP3R* was knocked down in all circadian neurons (Figure 3H). In the latter case, the coherence of peak phase was lost within groups (Rayleigh test, P > 0.1), consistent with the behavioral phenotypes of the two manipulations for *IP3R* RNAi. Knocking down *IP3R* in all circadian neurons caused stronger deficits in the amplitude of calcium rhythms of non-PDF-positive neurons (LNd, DN1, and DN3) than in those of PDF-positive neurons (s-LNv and l-LNv) (Figure S3A). The difference in the vulnerability to IP3R disruption between PDF-negative and PDF-positive neurons might explain why the *PDF-GAL4*-driven knockdown of *IP3R* affected neither calcium rhythms nor behavior. Together, these results implicate SERCA and IP3R channel activities as essential for slow calcium rhythms, and suggest the ER may be a key calcium source for the daily fluctuations of the basal calcium levels in circadian pacemaker neurons.

**Figure 3.**
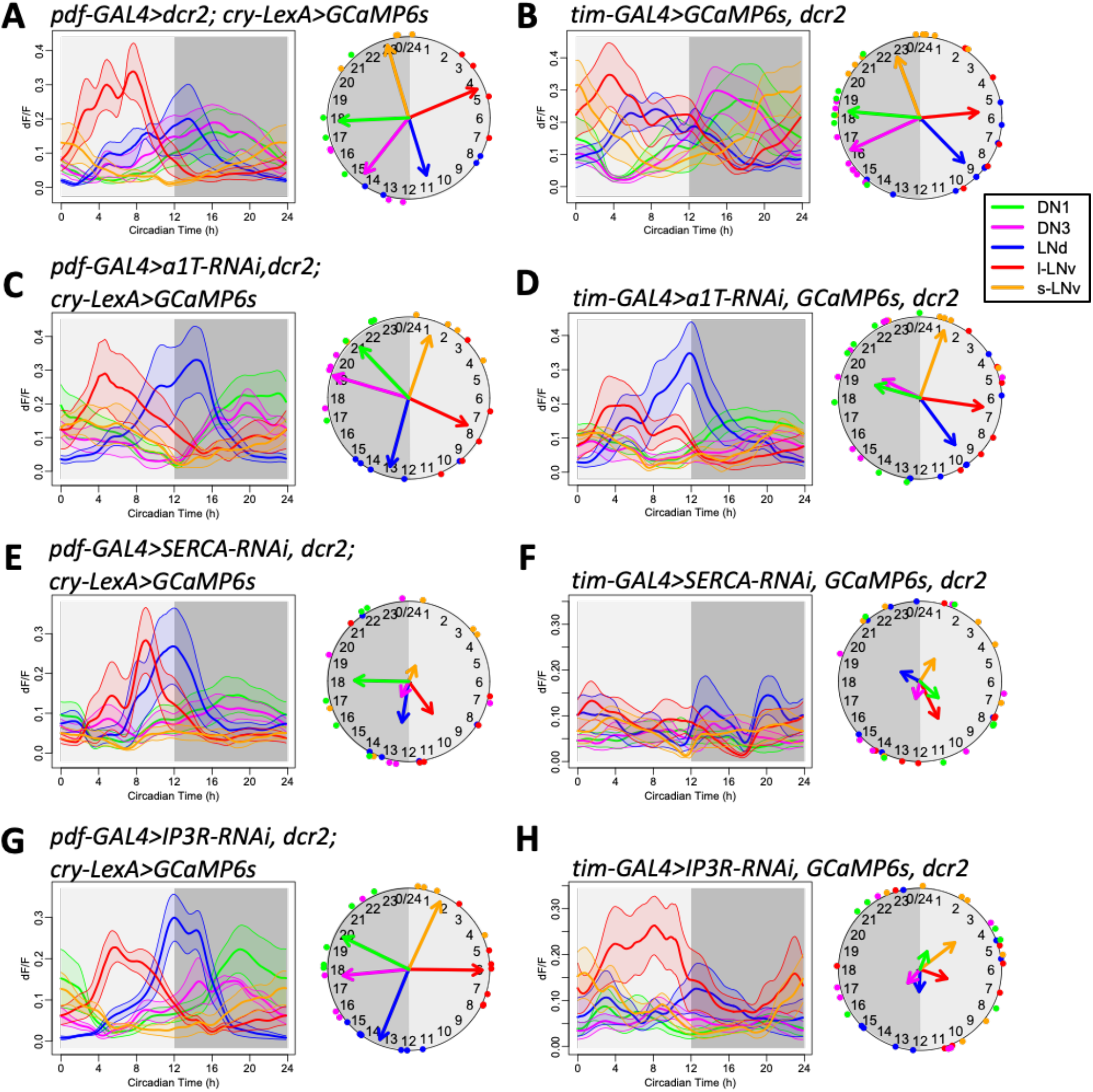
RNAi-screening for calcium channels required for circadian calcium rhythms. **(A-B)** Daily calcium activity rhythms of five major circadian pacemaker groups in control flies for (A) *pdf-GAL4*-driven knockdown (n = 4 flies) and (B) *tim-GAL4*-driven knockdown (n = 12 flies) in first day under constant darkness (DD1). **(C-D)** Daily calcium activity rhythms are normal in (C) *pdf-GAL4*-driven *a1T* knockdown (KK100082) flies (n = 5 flies) and (D) *tim-GAL4*-driven *a1T* knockdown flies (n = 7 flies). **(E-F)** Daily calcium activity rhythms are partially impaired in (E) *pdf-GAL4*-driven *SERCA* knockdown (KK107371) flies (n = 5 flies) and completely impaired in (F) *tim-GAL4*-driven *SERCA* knockdown flies (n = 6 flies). **(G-H)** Daily calcium activity rhythms are normal in (G) *pdf-GAL4*-driven *ip3r* knockdown flies (n = 7 flies) and impaired in (H) *tim-GAL4*-driven *ip3r* knockdown flies (n = 7 flies).

Because the slow calcium rhythms are driven by molecular clock gene oscillations (Liang et al 2016), we asked whether the molecular clock generates the slow calcium rhythms by regulating *SERCA* and *IP3R*. If *SERCA* and *IP3R* are downstream of the molecular clock, knocking down these genes would affect calcium rhythms and behavior but not affect the molecular clock itself. We examined PER protein levels in all five circadian pacemaker groups at four LD time points and found that PER cycling appeared robust in *IP3R*-knockdown flies, but was clearly diminished in *SERCA*-knockdown flies (Figure 4). Therefore, in this system, only *IP3R* appears to operate downstream of the molecular clock and is necessary to generate daily rhythms in basal calcium levels.

**Figure 4.**
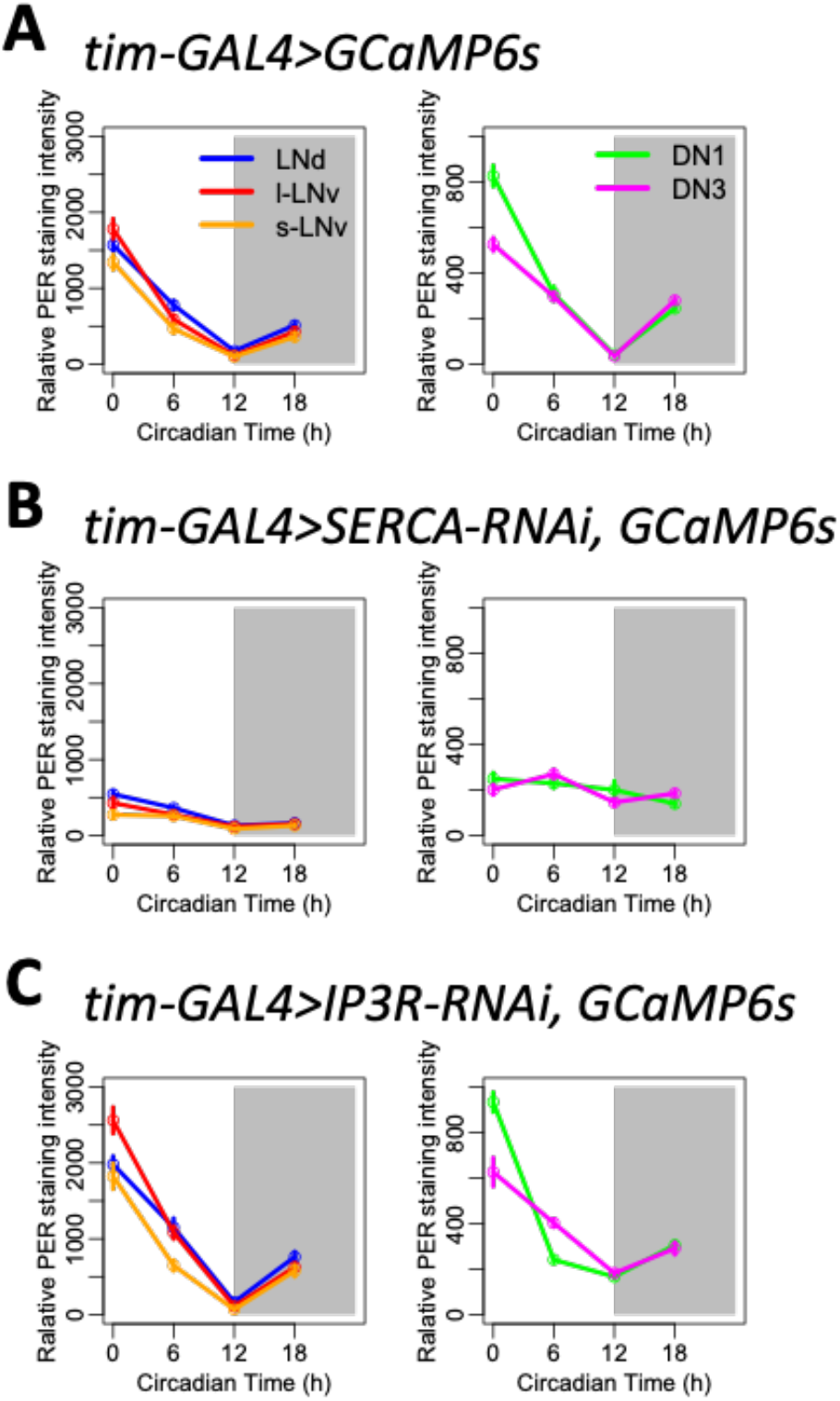
PER protein rhythms of control flies and flies with *SERCA* or *ip3r* knocked down in all pacemaker neurons. **(A)** Averaged PER protein staining intensity at four different time points (ZT0, ZT6, ZT12, and ZT18) in five groups of circadian pacemaker neurons from control flies. **(B)** PER protein rhythms are diminished when knocking down

### Fast calcium fluctuations require *α1T* calcium channels and *IP3R*

Knocking down the RNA for *α1T* voltage-gated calcium channels in pacemaker neurons impaired circadian rhythms in behavior, but did not affect circadian rhythmicity in basal calcium levels within those neurons. Therefore, we next asked whether *α1T* may underlie the circadian rhythm of fast calcium fluctuations in pacemakers. To test this, we again performed imaging across a series of short-term (2 min) high-frequency (1 Hz) calcium measurements on the same flies for a 24-hour day with 1-hour intervals (similar to Figure 1, yet with a slightly lower sampling rate and using flies that were WT for *cry*). By comparing control *Drosophila* to those with *α1T* knocked down in all circadian pacemaker neurons by *tim-GAL4*, we found that knocking down *α1T* did not affect the daily rhythms in the basal (slow) calcium level in any pacemaker group except for the DN1, which showed a reduction in the day-night difference of basal calcium level (Figure 5A-I and Figure S4). These high-frequency measures of slow basal calcium levels largely conform with those obtained with the slow-frequency (every 10 m) recording sessions (cf. Figure 3D). However, the high-frequency recording revealed that fast calcium fluctuations were significantly reduced in all circadian pacemaker neurons of the *α1T*-knockdown flies, except for the l-LNv, (Figure 5J and Figure S5). That specific pacemaker group instead displayed higher levels of fast calcium fluctuations. These results indicated that, at least in the majority of circadian pacemaker neurons, *α1T* is required for strong daily rhythms in fast calcium fluctuations and that the rhythm of fast calcium fluctuations can be selectively impaired.

**Figure 5.**
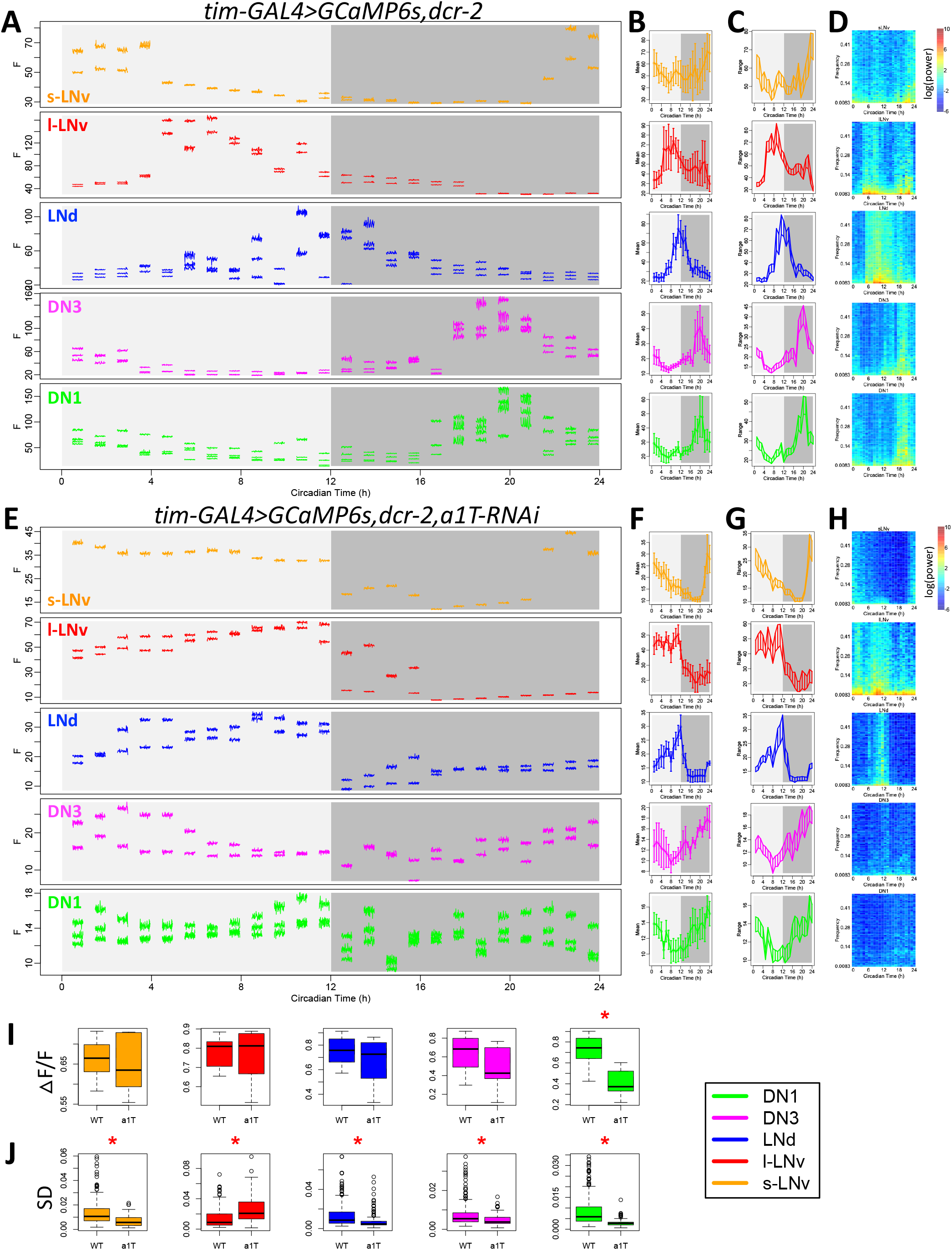
*a1T* knockdown reduces fast calcium fluctuations. **(A-D)** As Figure 1A-D, (A) raw calcium activity traces from one representative control fly. Each segmented trace is 2-min activity recorded at 1Hz. Averaged daily patterns of (B) mean calcium intensity, (C) the range of calcium transient, and (D) the power spectrum (n = 4 flies). **(E-H)** As in (A-D), raw calcium activity traces from one representative fly with *a1T* knockdown (KK100082) in all pacemaker neurons and averaged daily patterns of mean, range, and power spectrum in this genotype (n = 4 flies). **(I)** Box plot of daily range of calcium signal for individual neurons of five pacemaker groups between control flies and *a1T*-knockdown flies. The daily variation of DN1 calcium reduces in *a1T*-knockdown flies (t-test, *P < 0.05). **(J)** Box plot of standard deviations of calcium signal in each recording session for individual neurons of five pacemaker groups between control flies and *a1T*-knockdown flies. The standard deviations of fast calcium fluctuations in s-LNv, LNd, DN3, and DN1 are smaller, while that in l-LNv is larger in *a1T*-knockdown flies than that in control (t-test, *P < 0.05).

We then asked whether the fast calcium fluctuations are also affected in flies with *IP3R* knocked down in all pacemaker neurons. We performed the same high-frequency calcium imaging and analysis as in *α1T*-knockdown flies (Figure 6). The daily variation of pacemaker calcium was greatly reduced in *IP3R* -knockdown flies (Figure 6E), which largely recapitulated previous observations with less frequent sampling (cf. Figure 3H). In addition, fast calcium fluctuations were significantly reduced in all circadian pacemaker neurons of the *IP3R*-knockdown flies (Figure 6F and Figure S5). These results suggested that rhythms in basal calcium levels, regulated by *IP3R*, may be necessary for rhythms in fast calcium fluctuations.

**Figure 6.**
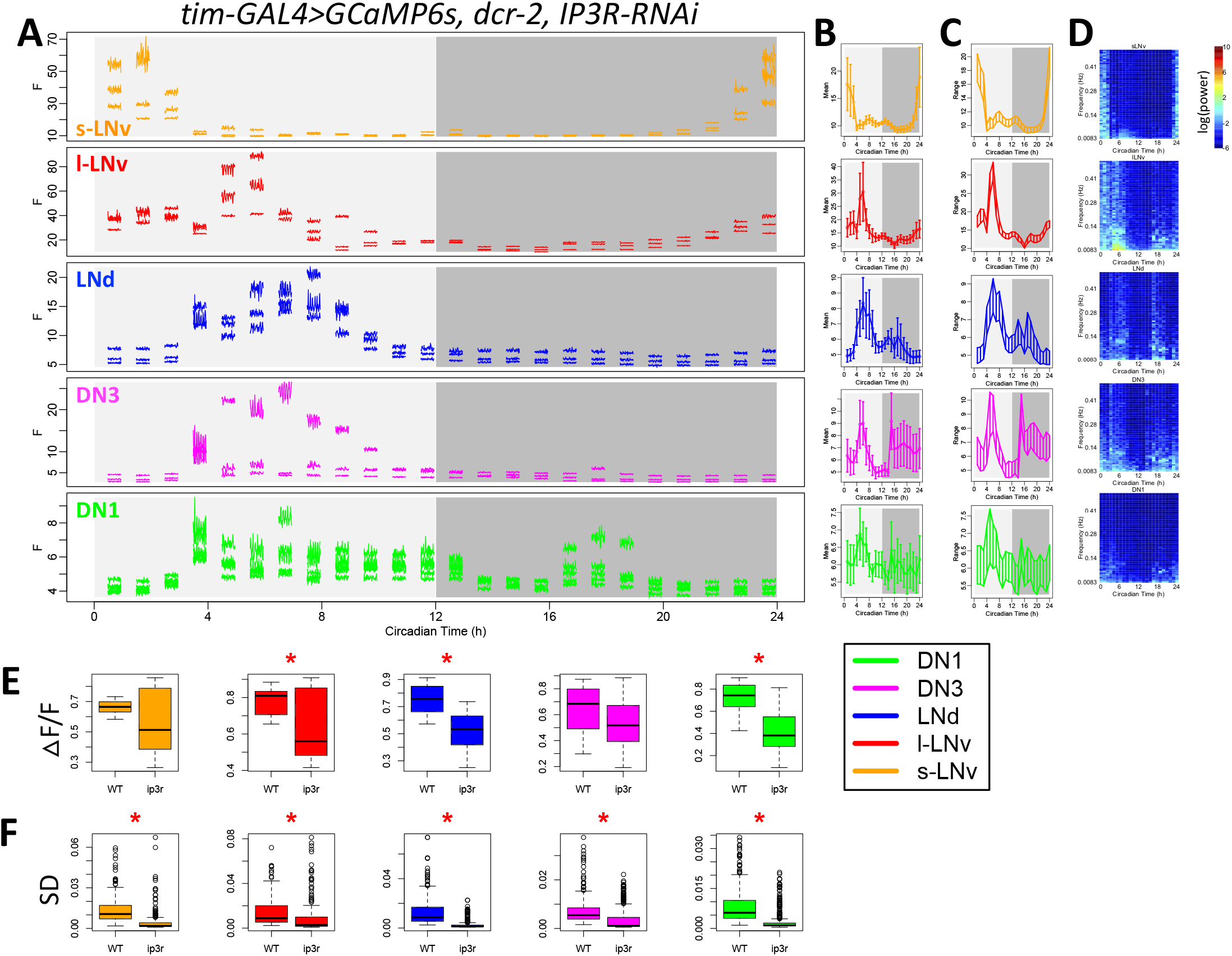
*IP3R* knockdown reduces fast calcium fluctuations. **(A-D)** As Figure 1A-D, (A) raw calcium activity traces from one representative fly with *a1T* knockdown in all pacemaker neurons. Each segmented trace is 2-min activity recorded at 1Hz. Averaged daily patterns of (B) mean calcium intensity, (C) the range of calcium transient, and (D) the power spectrum (n = 5 flies). **(E)** Box plot of daily range of calcium signal for individual neurons of five pacemaker groups between control flies and *IP3R*-knockdown flies. The daily variation of l-LNv, LNd, and DN1 calcium reduces in *IP3R*-knockdown flies (t-test, *P < 0.05). **(F)** Box plot of standard deviations of calcium signal in each recording session for individual neurons of five pacemaker groups between control flies and *IP3R*-knockdown flies. The standard deviations of fast calcium fluctuations in all circadian pacemaker neuron groups are smaller in *IP3R*-knockdown flies than that in control (t-test, *P < 0.05).

## Discussion

In this study, we used *in vivo* 24-hr high-frequency calcium imaging and genetic screening to study the cellular biology of daily calcium rhythms in circadian pacemaker neurons of *Drosophila*. We found that the calcium rhythm is in fact a composite: it reflects daily fluctuations in both a slow component (basal levels) and a fast one (high frequency fluctuations). We interpret fast calcium fluctuations as representations of calcium dynamics that occur as neurons fire single action potentials or bursts of them. While it is not sufficient to resolve single action potentials, GCaMP6-induced fluorescence is a good index of neuronal electrical activity (e.g. Chen et al., 2013; Greenberg et al 2018). For individual identified pacemakers, these two calcium rhythms share the same daily pattern, yet distinct calcium sources appear to contribute differentially to these two rhythms. An extracellular calcium influx, through plasma membrane calcium channels that include the *α1T* subunit, is critical for the fast calcium fluctuations. In contrast, calcium fluxes from the ER via the channel *IP3R* are required for both the slow rhythms in the basal calcium levels and the fast ones. Importantly, both channels are essential for normal circadian behavior. Thus, the molecular clocks may drive circadian rhythms in pacemaker neuron output by regulating different calcium sources to generate coordinate, but distinct rhythms in its calcium activities.

Circadian calcium rhythms (CCR) are widespread across taxa (Knight et al 1991; Colwell 2000). Calcium rhythms are required for circadian pacemaker functions in both rodents and *Drosophila* (Lundkvist et al 2005; Harrisingh et al 2007). Studies on mammalian circadian pacemakers in the suprachiasmatic nucleus (SCN) are controversial regarding the temporal relationship between the CCR and rhythms in electrical activity, such as in spontaneous firing rate (SFR) and in resting membrane potential (RMP). Recordings from SCN slice cultures showed that the phases of CCR in individual pacemakers are diverse and could be different from the populational phase of SFR rhythms (Ikeda et al 2003; Enoki et al 2012). However, the populational SFR phase is composed of many diverse phases of SFR rhythms on the individual cell level (Vanderleest et al, 2007); it is unclear whether SFR phases align with the phases of CCR. Imaging with both voltage sensor and calcium sensors in SCN slices, Brancaccio et al (2017) concluded that RMP rhythms and CCR were in phase, yet Enoki et al (2017) concluded that RMP rhythms and CCR were in phase in the ventral SCN, but in dorsal SCN, the CCR phase-led the RMP rhythms by about 2 hours. Because the voltage sensor signal measured from dorsal SCN may derive from the neural processes of ventral SCN neurons, the cellular interpretation of these results is unclear. Another source for the inconsistency might be the culture conditions: when SCN neurons were recorded *in vivo* by photometry, the rhythms in fast calcium activity were in phase with slow calcium rhythms (Jones et al 2018). In general, comparisons of population rhythms and rhythms in single cells are not easily reconciled. Our recordings that tracked pacemaker neurons from different identified groups *in vivo* for 24 hours showed that at the single cell level, slow calcium rhythms (CCR) were in phase with rhythms in fast calcium fluctuations; the latter likely reflect rhythms in SFR (Figure 1).

To measure the fast calcium fluctuations, we employed high-frequency light-sheet scanning with the wavelength of light that may activate pacemaker neurons and alter molecular clocks (Fogle et al 2011). We used *cry*^01^ flies to avoid the direct light responses of pacemaker neurons and found that all pacemaker groups displayed slow calcium rhythms, comparable to those we previously reported (Liang et al 2016), except for the l-LNv (Figure 1) which showed an additional calcium peak in the early evening. Because l-LNv innervate the optic lobes and receive large-scale visual inputs (Ashmore and Sehgal 2003), we speculate that the repeated optical scanning might activate l-LNv in the evening via visual systems. In the later experiments, when we reduced the illumination duration per hour from 31.5 sec (Figure 1, 7 ms exposure time, 15 frames per stack, 5Hz for 1min) to 2.4 sec (Figure 5 & 6, 1 ms exposure time, 20 frames per stack, 1Hz for 2 min), even in *cry* wild-type flies l-LNv did not show the evening peak. All pacemaker groups displayed slow calcium rhythms (Figure 5A), with the same phases as those obtained with the slow-frequency (every 10 min) recording sessions (cf. Figure 4D and Liang et al 2016). Therefore, by carefully tuning the illumination intensity for calcium imaging, we could monitor normal slow calcium rhythms and fast calcium fluctuations from the same individual pacemaker neurons.

The causal relationships between clock gene rhythms, calcium rhythms, and electrical activity rhythms in the SCN remain generally unresolved. Treating SCN slices with TTX (to block Na-dependent action potentials) diminished SFR rhythms (Ikeda et al 2003), partially affected RMP rhythms and CCRs (Hong et al 2012; Enoki et al 2012; Enoki et al 2017), and slowly affected clock gene rhythms over several days (Yamaguchi et al 2003). Dispersed SCN cells *in vitro* showed a TTX-resistant CCR, suggesting that CCR is driven by clock gene rhythms (Noguchi et al 2017). Thus, the variation in CCR sensitivity to TTX treatment might be caused by the degree to which clock gene rhythms *in vitro* become progressively dysfunctional. In *Drosophila*, our findings suggested that clock gene rhythms drive two components of the CCR -both basal calcium levels and fast calcium fluctuations -via circadian regulation of the ER channel *IP3R* and membrane voltage-gated calcium channel *α1T*. Both channels might then contribute to SFR and RMP rhythms. Similarly, in SCN pacemakers, pharmacologically blocking another ER channel *RyR* affected both CCR and SFR rhythms (Ikeda et al 2003), suggesting that rhythms in basal calcium levels are regulated by calcium from ER and are required for fast electric activity rhythms. In addition, SCN pacemakers also showed a circadian rhythm in fast calcium activity mediated by L-type voltage-gated calcium channels (Pennartz et al 2002). Pharmacologically blocking these membrane channels affected SFR rhythms and in some case affected CCR (Ikeda et al 2003; Enoki et al 2017). In our studies, manipulating a membrane voltage-gated calcium channel in all or a subset of pacemakers selectively affected rhythms in fast calcium fluctuations, which likely reflected SFR rhythms and thus impaired circadian outputs; however, it did not significantly affect the slow rhythms in basal calcium levels (Figure 5). Manipulating the ER calcium channel IP3R in all pacemakers affected rhythms in both slow and fast calcium rhythms. Therefore, in parallel to mammalian SCN neurons, *Drosophila* circadian pacemakers generate calcium rhythms by regulating both ER and extracellular calcium sources. Since our results suggest little or no role for the RyR channel (Figure 2B), the daily rhythmic regulation in fly pacemakers acts on a different set of ER and cytoplasmic membrane channels from those in mammalian pacemakers.

The RNAi knockdown experiments indicate a role for the ER calcium channel SERCA in supporting slow calcium rhythms in *Drosophila* pacemakers and behavioral rhythms. However, the high degree of lethality and the strong effects of SERCA-knockdown on the PER molecular oscillation precluded an assessment of its precise role. We conclude that *SERCA*, which maintains the ER-cytoplasmic calcium gradient, is essential for the normal physiology of the cells. In contrast to *SERCA*, our results support a hypothesis that *IP3R* is crucial actuator of the molecular clock. Previous transcriptomic analysis also supports that possibility: in circadian neurons, *IP3R* displays rhythmic expression, while *SERCA* does not (Figure S7; Abruzzi et al 2017).

Finally, the RNAi knockdown of plasma membrane calcium channel α1T indicated a role for voltage-gated T-type calcium channels in the final rhythmic output of the pacemakers. Consistent with a role in the presumed output pathway, impairing the rhythm of fast calcium activity strongly affected circadian behavior but did not affect the molecular clock or the slow calcium rhythms. T-type channels play a crucial role in other pacemakers such as the SA node of the mammalian heart (Vassort et al., 2006; Ono et al., 2010). Their conduction in the hyperpolarized state, and closure at more depolarized potentials, is central to their role in generating bursting dynamics with periods much longer than the membrane time constant (Perez-Reyes, 2003). Given the power spectrum of the fluctuations we observed, it seems possible they play a similar role in the “fast” (∼0.1Hz) fluctuations of Drosophila circadian neurons. Remarkably, the expression of *α1T* also displays a circadian rhythm, with distinct phases in different groups of pacemaker neurons (Figure S7; Abruzzi et al 2017). Collectively, these results suggest that *IP3R* and *α1T* channel activity are together critical to produce clock regulation of rhythms in both slow and fast calcium activities of critical pacemaker neurons.

## Methods

### Fly stocks

Flies were reared on standard yeast-supplemented cornmeal/agar food at room temperature. After eclosion, male flies were entrained under 12 h light: 12 h dark (LD) cycles at 25°C for at least 3 days. The *tim>GCaMPS6s; cry*^*01/01*^ flies were entrained under LD for more than 6 days. The following fly lines were previously described: *tim(UAS)-GAL4* (Blau & Young 1999), *pdf-GAL4* (Renn et al 1999), *cry-LexA* (Liang *et al*., 2017), *UAS-GCaMP6s* and *LexAop-GCaMP6s* (Chen *et al*., 2013). *UAS-dSTIM* and *UAS-dOrai* (Agrawal et al 2010) were gifts from Dr. G Hasan (NCBS, India). *UAS-ip3rRNAi* (Liu et al 2016) was a gift from Dr. M Wu (Johns Hopkins U.). Stable line: *UAS-dcr2; tim(UAS)-GAL4; UAS-GCaMP6s* and *pdf-GAL4;UAS-dcr2;cry-LexA,LexAop-GCaMP6s* were created for RNAi screening of calcium channels. The *cry-LexA* line was a gift from Dr. F Rouyer (CNRS Gyf, Paris).

RNAi lines were obtained from Bloomington Stock Center, Vienna Drosophila Resource Center, and Tokyo Stock Center. Two lines for *cac* (CG43368): *UAS-KK101478-RNAi* (VDRC 104168) and *UAS-JF02572-RNAi* (BDSC 27244). Two lines for *α1T* (CG15899): *UAS-KK100082-RNAi* (VDRC 108827) and *UAS-JF02150-RNAi* (BDSC 26251). Three lines for *α1D* (CG4894): *UAS-GD1737-RNAi* (VDRC 51491), *UAS-JF01848-RNAi* (BDSC 25830), and *UAS-HMS00294-RNAi* (BDSC 33413). Two lines for *α2δ* (CG12295): *UAS-KK101267-RNAi* (VDRC 18569) and *UAS-JF01825-RNAi* (BDSC 25807). One line for *Orai* (CG11430) *UAS-HMC03562-RNAi* (BDSC 53333). Three lines for *dSTIM* (CG9126): *UAS-KK102366-RNAi* (VDRC 106256), *UAS-GLC01785-RNAi* (BDSC 51685), and *UAS-JF02567-RNAi* (BDSC 27263). Two lines for *SERCA* (CG3725): *UAS-KK107371-RNAi* (VDRC 107446) and *UAS-JF01948-RNAi* (BDSC 25928). Three lines for *RyR* (CG19844): *UAS-KK101716-RNAi* (VDRC 109631), *UAS-HM05130-RNAi* (BDSC 28919), and *UAS-JF03381-RNAi* (BDSC 29445).

### *In vivo* fly preparations and calcium imaging

The fly surgery followed procedures previously described (Liang et al 2016; 2017). Flies were first anesthetized by CO_2_ and immobilized by inserting the neck into a narrow cut in an aluminum foil base. A portion of the dorso-anterior cuticle on one side of the head, an antenna, and a small part of one compound eye were then removed. For slow calcium rhythm measurements, imaging was conducted with a custom horizontal-scanning Objective Coupled Planar Illumination (hsOCPI) microscope (Holekamp *et al*., 2008). The scanning was done by moving the stage horizontally every 10 min for 24 hours. Each scan contained 20-40 separate frames with a step size of 5 to 10 microns. For fast calcium rhythm measurements, imaging was conducted with a custom high-speed dual-channel Objective Coupled Planar Illumination (OCPI-II) microscope (Greer and Holy 2019). Each scanning session involved moving the objective using a piezo motor at 1-5 Hz for 1-2 min. The same scans were then repeated on the same specimens every hour, for 24 hrs. During both slow and fast imaging modes, HL3 saline was continuously perfused (0.1-0.2 mL/min).

### Locomotor activity rhythm

Trikinetics *Drosophila* Activity Monitor (DAM) system was used to monitor the locomotor activity rhythms of individual flies. 4-6-day-old male flies were monitored for 6 days under light-dark (LD) cycles and then for 9 days under constant darkness (DD) condition. The circadian rhythmicity and periodicity were measured by χ^2^ periodogram with a 95% confidence cutoff and SNR analysis (Levine et al 2002). Arrhythmicity were defined by a power value (χ^2^ power at best period) less than 10, width lower than 1, a period less than 18 hours or more than 30 hours.

### Immunocytochemistry

The flies were entrained for 6 days under LD and dissected at ZT0, ZT6, ZT12, and ZT18. After dissected in ice-cold, calcium-free saline, fly brains were fixed for 15 min in 4% paraformaldehyde containing 7% picric acid (v/v) in PBS. Primary antibodies were rabbit anti-PER (1:5000; kindly provided by Dr. M. Rosbash, Brandeis Univ.; Stanewsky *et al*., 1997). Secondary antisera were Cy3-conjugated (1:1000; Jackson Immunoresearch, West Grove, PA). Images were taken on the Olympus FV1200 confocal microscope. PER protein immunostaining intensity was measured in ImageJ-based Fiji (Schindelin *et al*., 2012).

### Imaging data analysis

Calcium imaging data was acquired by a custom software, Imagine (Holekamp *et al*., 2008) and pre-processed using custom scripts in Julia 0.6 to produce non-rigid registration, alignment and maximal projection along z-axis. The images were then visualized and analyzed in ImageJ-based Fiji by manually selecting regions of interest (ROIs) over individual cells or groups of cells and measuring the intensity of ROIs over time. Slow calcium activity was analyzed as described previously (Liang *et al*., 2016, 2017). Fast calcium activity in each scanning session was analyzed similarly. Between sequential scanning sessions, the ROIs for individual neurons were manually corrected for position drifts. For the calcium signal of each ROI in each session, the mean of calcium intensity and the range of calcium intensity change was measured, and the power spectrum was generated by fast Fourier transform. Then the calcium signal was filtered by a high-pass filter (1/15 Hz) and the standard deviation of calcium changes was measured. Calcium activity trace analysis and statistics were performed using R 3.3.3 and Prism 8 (GraphPad, San Diego CA).

## Acknowledgments

We thank Cody Greer for building the OCPI-2 microscope, the Holy and Taghert laboratories for advice, and the Washington University Center for Cellular Imaging (WUCCI) for technical support. Gaiti Hasan, Mark Wu, Francois Rouyer, Michael Rosbash, Bloomington Stock Center, Vienna Drosophila Resource Center, and Tokyo Stock Center provided fly stocks and reagents. The work was supported by the Washington University McDonnell Center for Cellular and Molecular Neurobiology and by NIH grants R01 NS068409 and R01 DP1 DA035081 (T.E.H.), R01 NS099332 and R01 GM127508 (P.H.T.), and R24 NS086741 (T.E.H. and P.H.T.).

## Author contributions

X.L., T.H.E., and P.H.T. conceived the experiments; X.L. performed and analyzed all experiments; X.L., P.H.T. and T.H.E. wrote the manuscript.

## References

Abruzzi, K.C., Zadina, A., Luo, W., Wiyanto, E., Rahman, R., Guo, F., Shafer, O., and Rosbash, M. (2017). RNA-seq analysis of Drosophila clock and non-clock neurons reveals neuron-specific cycling and novel candidate neuropeptides. PLOS Genet. 13, e1006613.

Agrawal, N., Venkiteswaran, G., Sadaf, S., Padmanabhan, N., Banerjee, S., & Hasan, G. (2010). Inositol 1, 4, 5-trisphosphate receptor and dSTIM function in Drosophila insulin-producing neurons regulates systemic intracellular calcium homeostasis and flight. Journal of Neuroscience, 30(4), 1301–1313.

Ashmore, L.J., and Sehgal, A. (2003). A Fly’s Eye View of Circadian Entrainment. J. Biol. Rhythms 18, 206–216.

Berridge, M.J. (1998). Neuronal calcium signaling. Neuron 21, 13–26.

Blau, J., and Young, M.W. (1999). Cycling vrille expression is required for a functional Drosophila clock. Cell 99, 661–671.

Brancaccio, Marco, Elizabeth S. Maywood, Johanna E. Chesham, Andrew SI Loudon, and Michael H. Hastings. “A Gq-Ca2+ axis controls circuit-level encoding of circadian time in the suprachiasmatic nucleus.” Neuron 78, no. 4 (2013): 714–728.

Cao, G., & Nitabach, M. N. (2008). Circadian control of membrane excitability in Drosophila melanogaster lateral ventral clock neurons. Journal of Neuroscience, 28(25), 6493–6501.

Chen, T.-W., Wardill, T.J., Sun, Y., Pulver, S.R., Renninger, S.L., Baohan, A., Schreiter, E.R., Kerr, R. a, Orger, M.B., Jayaraman, V., et al. (2013). Ultrasensitive fluorescent proteins for imaging neuronal activity. Nature 499, 295–300.

Chorna, T., and Hasan, G. (2012). The genetics of calcium signaling in Drosophila melanogaster. Biochim. Biophys. Acta - Gen. Subj. 1820, 1269–1282.

Colwell, C.S. (2000). Circadian modulation of calcium levels in cells in the suprachiasmatic nucleus. Eur. J. Neurosci. 12, 571–576.

Colwell, C.S. (2011). Linking neural activity and molecular oscillations in the SCN. Nat. Rev. Neurosci. 12, 553–569.

Dunlap, J. (1999). Molecular bases for circadian clocks. Cell, 96, 271–290.

Enoki, R., Kuroda, S., Ono, D., Hasan, M.T., Ueda, T., Honma, S., Honma, K., and Sens, H. (2012). Topological specificity and hierarchical network of the circadian calcium rhythm in the suprachiasmatic nucleus. Proc. Natl. Acad. Sci. U. S. A. 109, 21498–21503.

Enoki, R., Oda, Y., Mieda, M., Ono, D., Honma, S., and Honma, K.-I. (2017). Synchronous circadian voltage rhythms with asynchronous calcium rhythms in the suprachiasmatic nucleus. Proc. Natl. Acad. Sci. 114, E2476–E2485.

Flourakis, M., Kula-Eversole, E., Hutchison, A.L.L., Han, T.H.H., Aranda, K., Moose, D.L.L., White, K.P.P., Dinner, A.R.R., Lear, B.C.C., Ren, D., et al. (2015). A Conserved Bicycle Model for Circadian Clock Control of Membrane Excitability. Cell 162, 836–848.

Fogle, K.J., Parson, K.G., Dahm, N. a, and Holmes, T.C. (2011). CRYPTOCHROME is a blue-light sensor that regulates neuronal firing rate. Science 331, 1409–1413.

Greer, C. J., & Holy, T. E. (2019). Fast objective coupled planar illumination microscopy. Nature communications, 10(1), 1–14.

Greenberg, D. S., Wallace, D. J., Voit, K. M., Wuertenberger, S., Czubayko, U., Monsees, A., … & Kerr, J. N. (2018). Accurate action potential inference from a calcium sensor protein through biophysical modeling. BioRxiv, 479055.

Harrisingh, M.C., Wu, Y., Lnenicka, G. a, and Nitabach, M.N. (2007). Intracellular Ca2+ regulates free-running circadian clock oscillation in vivo. J. Neurosci. 27, 12489–12499.

Herzog, E.D. (2007). Neurons and networks in daily rhythms. Nat. Rev. Neurosci. 8, 790–802.

Holekamp, T.F., Turaga, D., and Holy, T.E. (2008). Fast three-dimensional fluorescence imaging of activity in neural populations by objective-coupled planar illumination microscopy. Neuron 57, 661–672.

Hong, J.H., Jeong, B., Min, C.H., and Lee, K.J. (2012). Circadian waves of cytosolic calcium concentration and long-range network connections in rat suprachiasmatic nucleus. Eur. J. Neurosci. 35, 1417–1425.

Ikeda, M., Sugiyama, T., Wallace, C.S., Gompf, H.S., Yoshioka, T., Miyawaki, A., and Allen, C.N. (2003). Circadian dynamics of cytosolic and nuclear Ca2+ in single suprachiasmatic nucleus neurons. Neuron 38, 253–263.

Itri, J.N., Michel, S., Vansteensel, M.J., Meijer, J.H., and Colwell, C.S. (2005). Fast delayed rectifier potassium current is required for circadian neural activity. Nat. Neurosci. 8, 650– 656.

Jones, J. R., Simon, T., Lones, L., & Herzog, E. D. (2018). SCN VIP neurons are essential for normal light-mediated resetting of the circadian system. Journal of Neuroscience, 38(37), 7986–7995.

Knight, M.R., Campbell, A.K., Smith, S.M., and Trewavas, A.J. (1991). Transgenic plant aequorin reports the effects of touch and cold-shock and elicitors on cytoplasmic calcium. Nature 352, 524–526.

Levine, J., Funes, P., Dowse, H., and Hall, J. (2002). Signal analysis of behavioral and molecular cycles. BMC Neurosci. 25, 1–25.

Liang, X., Holy, T. E., & Taghert, P. H. (2016). Synchronous Drosophila circadian pacemakers display nonsynchronous Ca2+ rhythms in vivo. Science, 351(6276), 976–981.

Liang, X., Holy, T. E., & Taghert, P. H. (2017). A series of suppressive signals within the Drosophila circadian neural circuit generates sequential daily outputs. Neuron, 94(6), 1173–1189.

Lundkvist, G.B., Kwak, Y., Davis, E.K., Tei, H., and Block, G.D. (2005). A calcium flux is required for circadian rhythm generation in mammalian pacemaker neurons. J. Neurosci. 25, 7682–7686.

Meredith, A.L., Wiler, S.W., Miller, B.H., Takahashi, J.S., Fodor, A. a, Ruby, N.F., and Aldrich, R.W. (2006). BK calcium-activated potassium channels regulate circadian behavioral rhythms and pacemaker output. Nat. Neurosci. 9, 1041–1049.

Mohawk, J. A, Green, C.B., and Takahashi, J.S. (2012). Central and peripheral circadian clocks in mammals. Annu. Rev. Neurosci. 35, 445–462.

Noguchi, T., Leise, T.L., Kingsbury, N., Diemer, T., Wang, L.L., Henson, M.A., and Welsh, . (2017). Calcium Circadian Rhythmicity in the Suprachiasmatic Nucleus: Cell Autonomy and Network Modulation. Eneuro ENEURO.0160-17.2017.

Ono K, and Iijima T. Cardiac T-type Ca(2+) channels in the heart. J. Mol. Cell. Cardiol. 2010 48, 65–70.

Panda, S., Antoch, M. P., Miller, B. H., Su, A. I., Schook, A. B., Straume, M., … & Hogenesch, J. B. (2002). Coordinated transcription of key pathways in the mouse by the circadian clock. Cell, 109(3), 307–320.

Pennartz, C.M. a, de Jeu, M.T.G., Bos, N.P.A., Schaap, J., and Geurtsen, A.M.S. (2002). Diurnal modulation of pacemaker potentials and calcium current in the mammalian circadian clock. Nature 416, 286–290.

Perez-Reyes E. Molecular Physiology of Low-Voltage-Activated T-type Calcium Channels Phys. Rev. (2003) 83: 117–161.

Pitts, G.R., Ohta, H., and McMahon, D.G. (2006). Daily rhythmicity of large-conductance Ca2+-activated K+ currents in suprachiasmatic nucleus neurons. Brain Res. 1071, 54–62.

Pologruto, T. A., Yasuda, R., & Svoboda, K. (2004). Monitoring neural activity and [Ca2+] with genetically encoded Ca2+ indicators. Journal of Neuroscience, 24(43), 9572–9579.

Renn, S.C., Park, J.H., Rosbash, M., Hall, J.C., and Taghert, P.H. (1999). A pdf neuropeptide gene mutation and ablation of PDF neurons each cause severe abnormalities of behavioral circadian rhythms in Drosophila. Cell 99, 791–802.

Schindelin, J., Arganda-Carreras, I., Frise, E., Kaynig, V., Longair, M., Pietzsch, T., Preibisch, S., Rueden, C., Saalfeld, S., Schmid, B., et al. (2012). Fiji: an open-source platform for biological-image analysis. Nat. Methods 9, 676–682.

Sheeba, V., Sharma, V.K., Gu, H., Chou, Y.-T., O’Dowd, D.K., and Holmes, T.C. (2008). Pigment dispersing factor-dependent and -independent circadian locomotor behavioral rhythms. J. Neurosci. 28, 217–227.

Stanewsky, R., Frisch, B., Brandes, C., Hamblen-Coyle, M. J., Rosbash, M., & Hall, J. C. (1997). Temporal and Spatial Expression Patterns of Transgenes Containing Increasing Amounts of the Drosophila Clock Gene period and a lacZ Reporter: Mapping Elements of the PER Protein Involved in Circadian Cycling. Journal of Neuroscience, 17(2), 676–696.

Streit AK, Fan YN, Masullo L, Baines RA. Calcium Imaging of Neuronal Activity in Drosophila Can Identify Anticonvulsive Compounds. PLoS One. 2016 Feb 10;11(2):e0148461. doi:10.1371/journal.pone.0148461. PubMed PMID: 26863447; PubMed Central PMCID: PMC4749298.

VanderLeest HT, Houben T, Michel S, Deboer T, Albus H, Vansteensel MJ, Block GD, Meijer JH. Seasonal encoding by the circadian pacemaker of the SCN. Curr Biol. 2007 Mar 6;17(5):468–73. PubMed PMID: 17320387.

Vassort G, Talavera K, and Alvarez JL. Role of T-type Ca2+ channels in the heart. Cell Calcium. (2006) 40, 205–20.

Welsh, D.K., Logothetis, D.E., Meister, M., and Reppert, S.M. (1995). Individual neurons dissociated from rat suprachiasmatic nucleus express independently phased circadian firing rhythms. Neuron 14, 697–706.

Welsh, D.K., Takahashi, J.S., and Kay, S. a (2010). Suprachiasmatic nucleus: cell autonomy and network properties. Annu. Rev. Physiol. 72, 551–577.

Yaksi, E., & Friedrich, R. W. (2006). Reconstruction of firing rate changes across neuronal populations by temporally deconvolved Ca 2+ imaging. Nature methods, 3(5), 377.

Yamaguchi, S., Isejima, H., Matsuo, T., Okura, R., Yagita, K., Kobayashi, M., and Okamura, H. (2003). Synchronization of Cellular Clocks in the Suprachiasmatic Nucleus. Science (80-.). 302, 1408–1412.

