## Supplemental figures for "Circadian pacemaker neurons display co-phasic rhythms in basal calcium level and in fast calcium fluctuations"

### Supplemental Materials

#### Figure S1

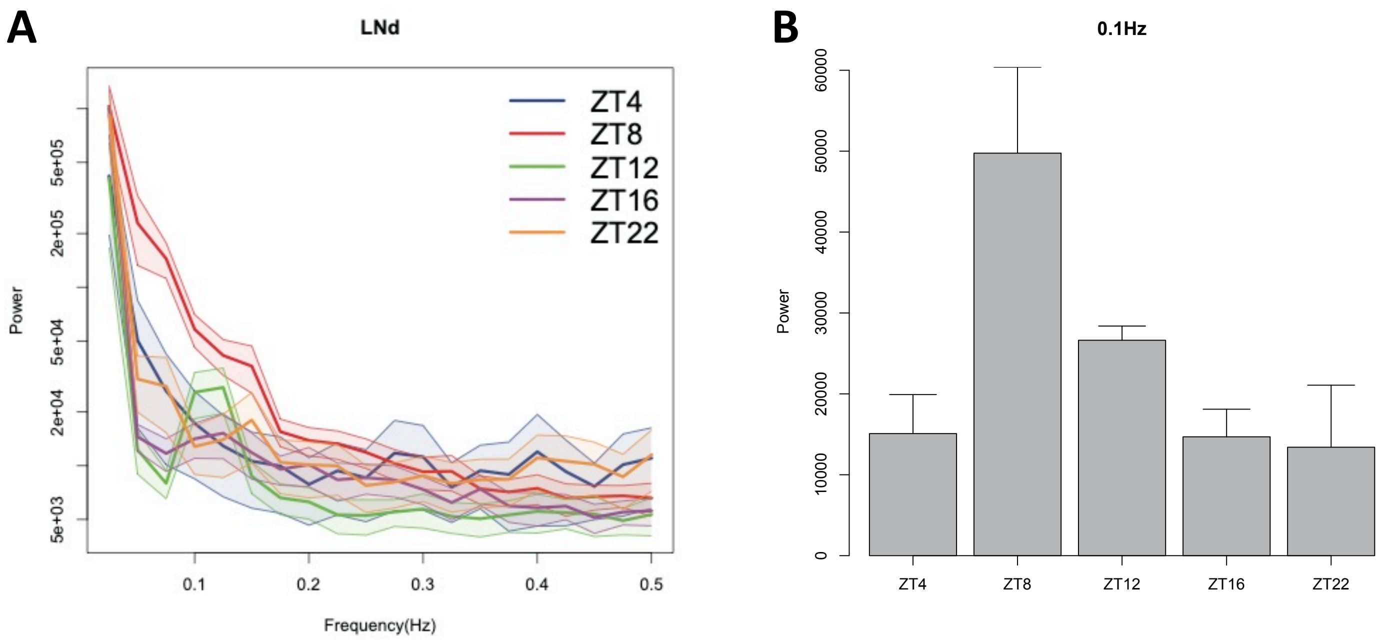


**Figure S1. Daily pattern of fast calcium activity in LNd.**
**(A)** Averaged power spectrums for five different timepoints (n > 6 flies for each timepoint). **(B)** Averaged power at 0.1Hz for five different timepoints shows a similar daily pattern as LNd slow calcium rhythms.

### Figure S2

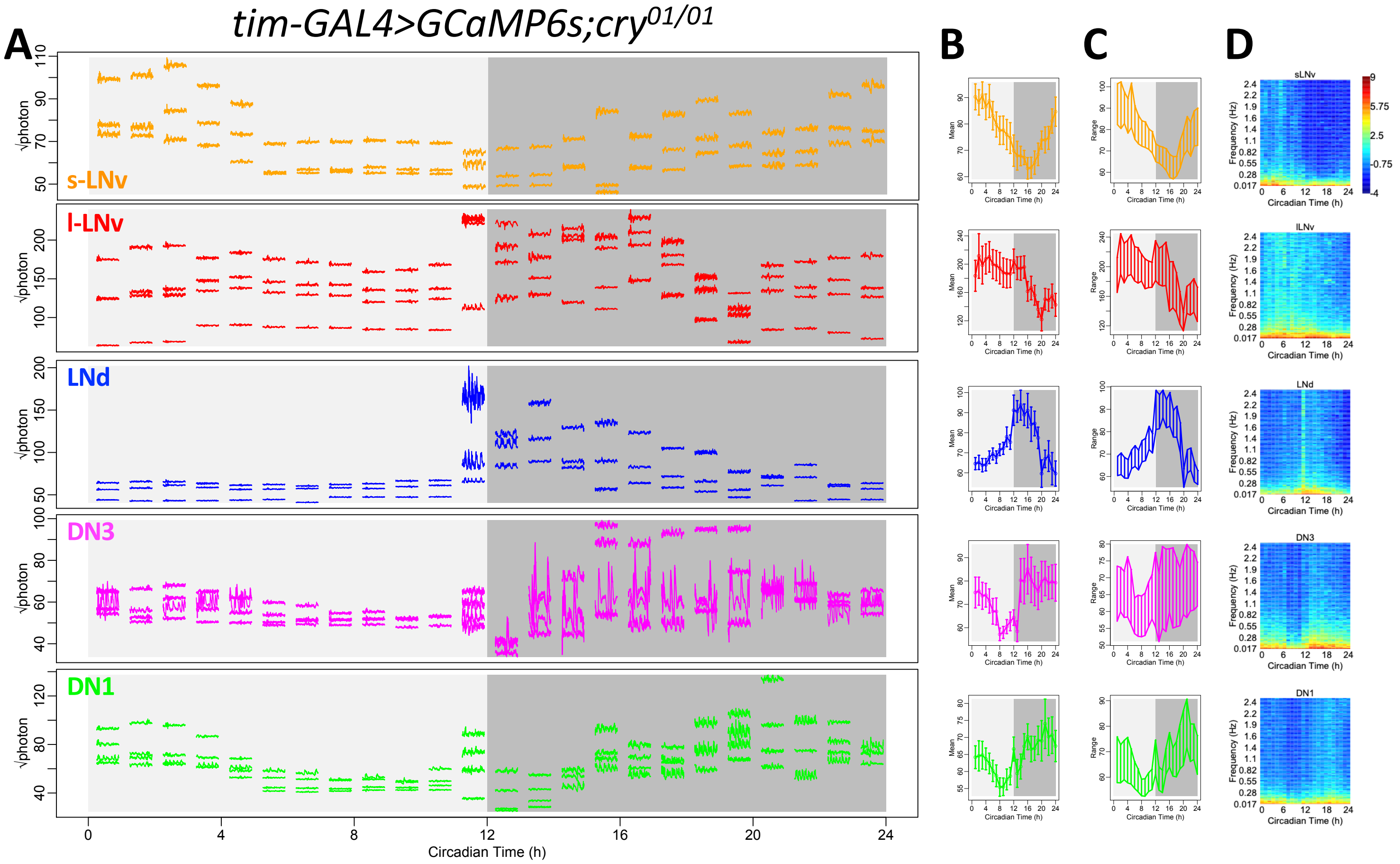


**Figure S2. Daily pattern of fast calcium activity in circadian pacemaker neurons normalized for shot noise.** **(A)** Raw calcium activity traces from the representative fly shown in Figure 1A. The intensity of calcium signal was calculated as the square root of photon number collected from individual region of interest (ROI), in order to normalize the effect of shot noise. **(B)** The mean of the photon number square root over the 1-min recording session at each of the 24 timepoints, averaged across all 6 flies studied. Error bars denote SEM. **(C)** Daily patterns of the range of calcium transients based on the square root of photon number collected from individual ROI, over the 1-min recording session at each of the 24 timepoints, averaged across all 6 flies studied. **(D)** Daily pattern of the power spectrum based on the square root of photon number collected from individual ROI, over the 1-min recording session at each of 24 timepoints, averaged across all 6 flies studied.

### Figure S3

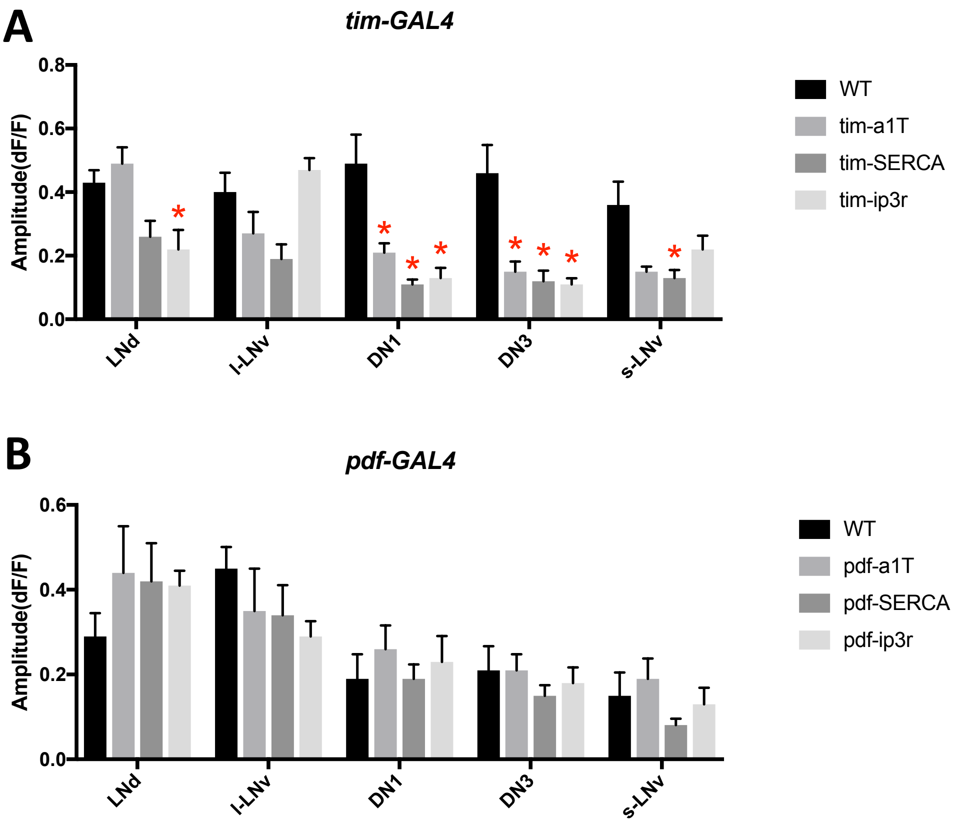


**Figure S3. Amplitude of daily calcium peaks in calcium-channel-knockdown flies.**
**(A)** The averaged amplitude of daily calcium peaks of five pacemaker groups in flies measured in Figure 3B, 3D, 3F, and 3H, wild type (*tim-GAL4>dcr2*) or with calcium channels knockdown by *tim-GAL4* (two-way ANOVA followed by Dunnett's multiple comparisons test, *P < 0.05). **(B)** The averaged amplitude of daily calcium peaks of five pacemaker groups in flies measured in Figure 3A, 3C, 3E, and 3G, wild type (*pdf-GAL4>dcr2*) or with calcium channels knockdown by *pdf-GAL4*.

Figure S4

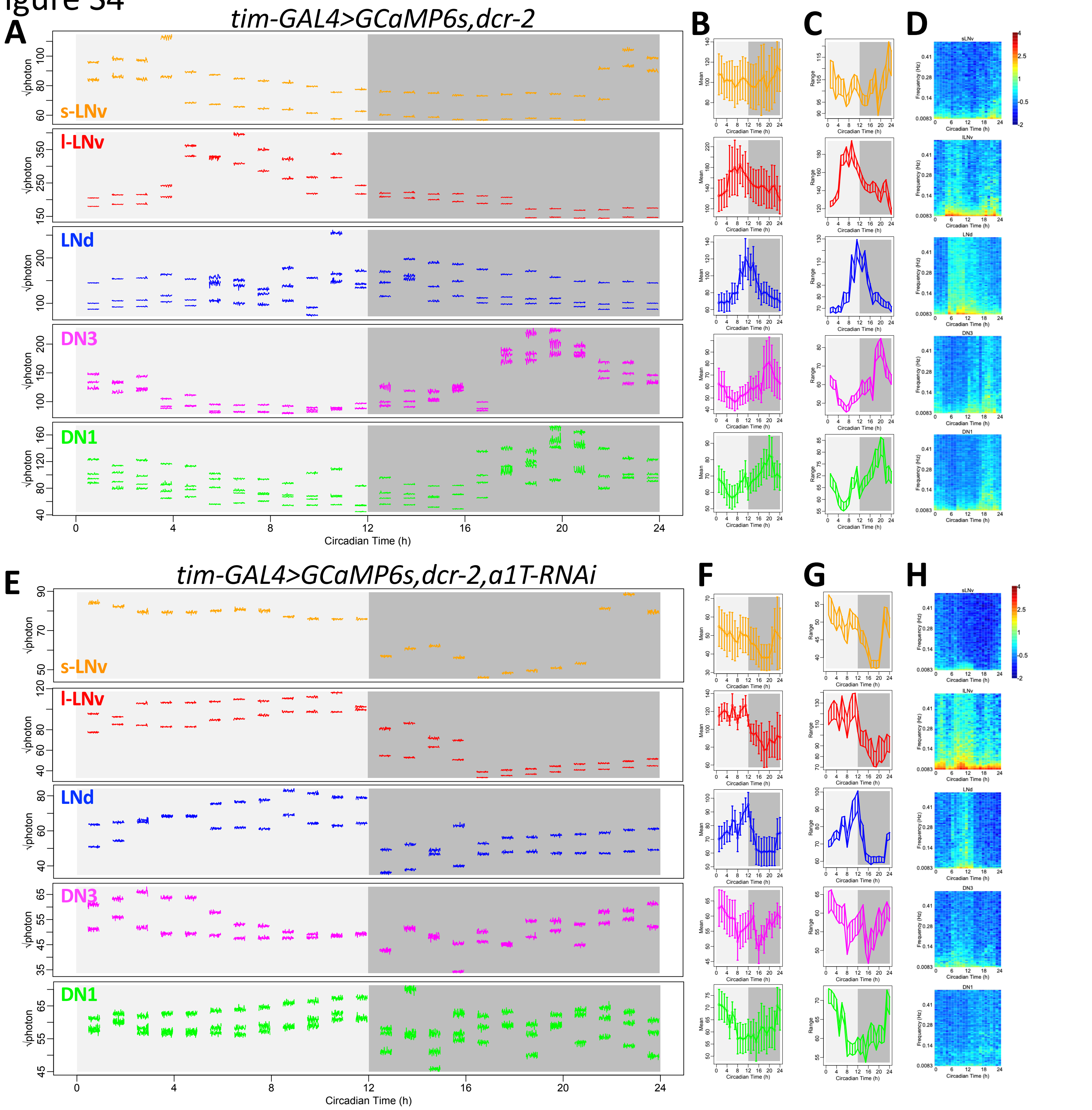


**Figure S4. Changes in shot noise levels do not explain reductions in fast calcium fluctuations by *a1T* knockdown.**

**(A-D)** As Figure S2A-D, (A) raw calcium activity traces from the representative fly shown in Figure 5A. The intensity of calcium signal was calculated as the square root of photon number collected from individual region of interest (ROI), in order to normalize the effect of shot noise. Averaged daily patterns of (B) mean calcium intensity, (C) the range of calcium transient, and (D) the power spectra were calculated based on the square root of photon number collected from individual ROIs (n = 4 flies). **(E-H)** As in (A-D), raw calcium activity traces from one representative fly shown in Figure 5E with *a1T* knockdown in all pacemaker neurons and averaged daily patterns of mean, range, and power spectrum in this genotype calculated based on the square root of photon number collected from individual ROIs (n = 4 flies).

### Figure S5

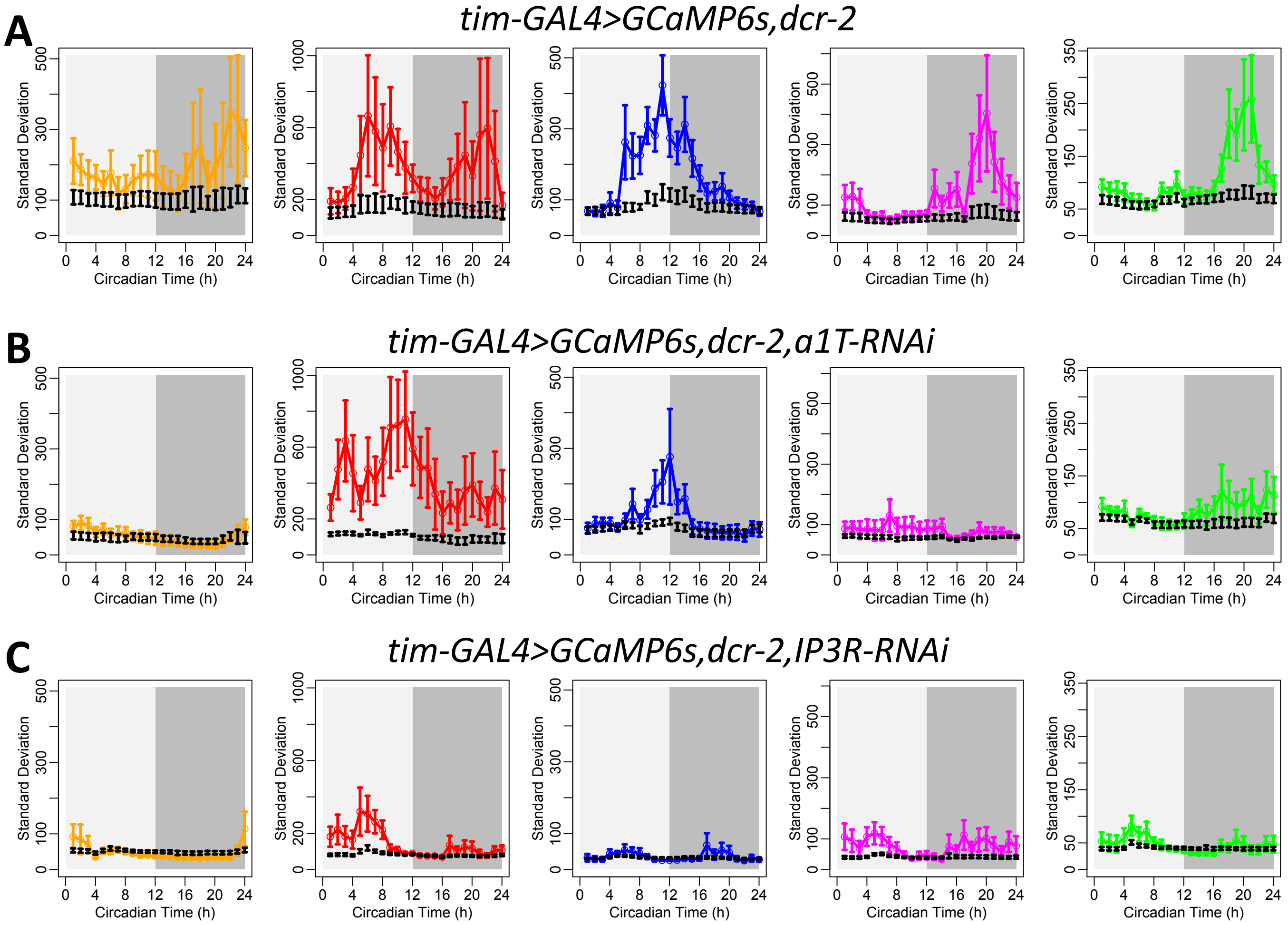


**Figure S5. Fast calcium fluctuations of *a1T* or *IP3R* knockdown were below the range of shot noise.** **(A)** Comparison of standard deviations between calcium activity and shot noise in wild type flies. Color lines, daily pattern of the standard deviations of calcium intensity (calculated as the collected photon number from individual ROIs) over the 1-min recording session at each of the 24 timepoints, averaged across all wide-type 4 flies studied in Figure 5A-D. Error bars denote SEM. For each timepoint, black error bars were the standard deviations of shot noise estimated by the square root of photon number collected from individual ROIs. At those time of day when mean calcium signal was high (shown in Figure 5B), the standard deviations of calcium signal were significantly higher than the standard deviations generated by shot noise , suggesting that the calcium fluctuation was generated by physiologically-relevant events. **(B)** Comparison of standard deviations between calcium activity and shot noise in flies shown in Figure 5E-H with *a1T* knockdown. Except for LNd and l-LNv, the standard deviations of calcium signal were not significantly different from the standard deviations generated by shot noise at any time of day. **(C)** Comparison of standard deviations between calcium activity and shot noise in flies shown in Figure 6 with *IP3R* knockdown.

### Figure S6

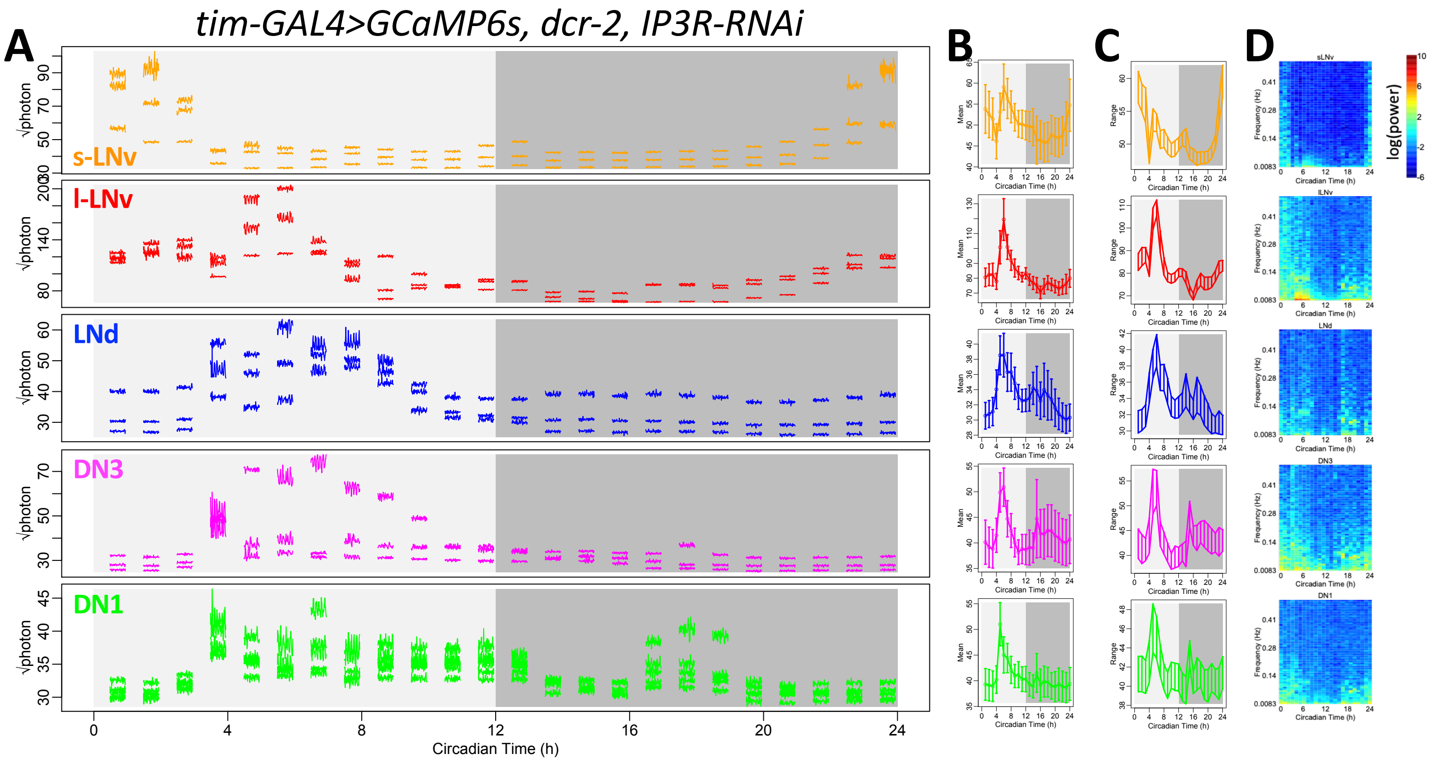


**Figure S6. Changes in shot noise levels do not explain reductions in fast calcium fluctuations by *IP3R* knockdown.**

**(A-D)** As Figure S2A-D, (A) raw calcium activity traces from the representative fly shown in Figure 6A. The intensity of calcium signal was calculated as the square root of photon number collected from individual region of interest (ROI), in order to normalize the effect of shot noise. Averaged daily patterns of (B) mean calcium intensity, (C) the range of calcium transient, and (D) the power spectrum was calculated based on the square root of photon number collected from individual ROIs (n = 5 flies).

### Figure S7

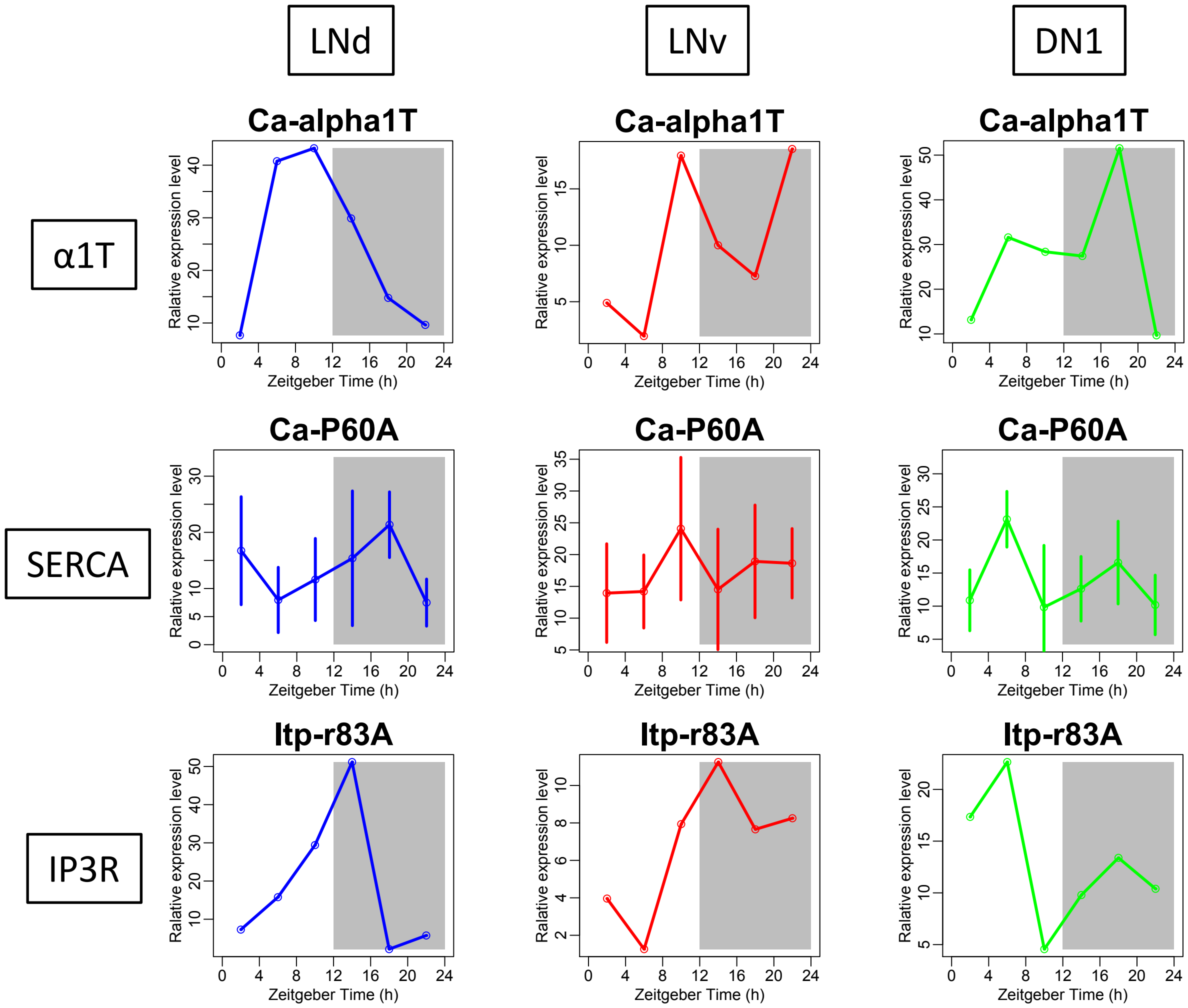


**Figure S7. Daily rhythms in expression level of different calcium channels.**
The daily expression pattern of *a1T*, *SERCA,* and *ip3r* in three different pacemaker groups LNd, PDF-positive LNv neurons, and DN1. Data is obtained from Abruzzi et al 2017, in which each pacemaker group was sorted and analyzed by RNA sequencing for two continuous days with 4-hour intervals.
